# *Disrupted-in-Schizophrenia-1* is required for proper pyramidal cell-interneuron communication and network dynamics in the prefrontal cortex

**DOI:** 10.1101/2021.12.10.472099

**Authors:** Jonas-Frederic Sauer, Marlene Bartos

## Abstract

We interrogated prefrontal circuit function in mice lacking *Disrupted-in-schizophrenia-1* (Disc1-mutant mice), a risk factor for psychiatric disorders. Single-unit recordings in awake mice revealed reduced average firing rates of fast-spiking interneurons (INTs), including optogenetically identified parvalbumin-positive cells, and a lower proportion of INTs phase-coupled to ongoing gamma oscillations. Moreover, we observed decreased spike transmission efficacy at local pyramidal cell (PYR)-INT connections *in vivo*, suggesting a reduced excitatory effect of local glutamatergic inputs as a potential mechanism of lower INT rates. On the network level, impaired INT function resulted in altered activation of PYR assemblies: While assembly activations were observed equally often, the expression strength of individual assembly patterns was significantly higher in Disc1-mutant mice. Our data thus reveal a role of Disc1 in shaping the properties of prefrontal assembly patterns by setting prefrontal INT responsiveness to glutamatergic drive.

## Introduction

Cognitive impairment is a major burden for individuals suffering from psychiatric disease. A series of investigation has put emphasis on the critical role of GABAergic INTs in the emergence of cognitive dysfunction in psychiatric illnesses (Akbarian et al., 1995, Mirnics et al., 2000, Volk et al., 2000, Lewis et al., 2005, Uhlhaas and Singer, 2010, Lisman, 2012, Volk et al., 2016, Ferguson and Gao, 2018). Among the various neocortical INT types, especially fast-spiking parvalbumin-positive interneurons (PVIs) have attracted substantial attention. Post mortem data indicated reduced numbers of PVIs and diminished expression of the GABA-synthetizing enzyme GAD in PVIs of the prefrontal cortex of schizophrenia patients (Lewis et al., 2005, Volk et al., 2016, Marín, 2012). Furthermore, eliminating prefrontal PVI output signaling induced working memory deficits in mice, underscoring the central role of PVI activity in cognitive function (Murray et al., 2015).

Transgenic Disc1-mutant mice, which resemble an ultra-rare mutation with high penetrance for schizophrenia and major depressive disorder (Blackwood et al., 2001), show cognitive deficits, thus rendering Disc1-mutant mice a prime candidate to investigate circuit mechanisms of a psychiatric disease (Koike et al., 2006, Kvajo et al., 2008). Much research has focused on the medial prefrontal cortex (mPFC) as an important center for cognitive function (Goldman-Rakic, 1995, Sigurdsson et al., 2010, Cho et al., 2015, Sauer et al., 2015, Kim et al., 2016a, Chini et al., 2020, Duvarci et al., 2018, Kaefer et al., 2020) but the circuit alterations underlying cognitive impairment are not well understood. *In vitro* studies in Disc1-mutants have put forward evidences for altered GABAergic and glutamatergic signaling (Sauer et al., 2015, Crabtree et al., 2017), but studies investigating the activity profile of prefrontal circuit components *in vivo* are lacking. Furthermore, whether impairments of PVIs might contribute to altered network function and cognitive deficits in Disc1-mutant mice is not known.

Here, we interrogated PVI function in the mPFC of Disc1-mutant mice. In awake animals, fast-spiking INTs and optogenetically identified PVIs showed reduced firing levels. Furthermore, the proportion of INTs phase-coupled to gamma (40-100 Hz) oscillations was markedly reduced. Using spike train crosscorrelation, we demonstrate that the excitatory effect of local glutamatergic synaptic inputs is reduced *in vivo*, consistent with impaired excitability of PVIs. On the network level, these alterations in INT function impacted the activation of cortical cell assemblies, which are thought to be the building blocks of cortical information processing (Buzsáki, 2010).

## Results

### Impaired interneuron activity in the medial prefrontal cortex of Disc1-mice

Disc1-mutant mice have been shown to express impaired execution of working memory (Koike et al., 2006, Kvajo et al., 2008). We confirmed working memory deficits in this mouse line in a delayed non-match-to place paradigm (**Supplementary Fig. 1**). To assess the potential network alterations underlying cognitive impairment, we performed single-unit recordings from the mPFC of Disc1-mutant and control mice that were awake in their home cage (n=1443 units) and analyzed discharge rates of PYRs and INTs (**Fig. 1**). Both neuron types were separated based on spike waveform kinetics (Sirota et al., 2008, PYRs n=1265, putative INTs n=178; 9 Disc1 and 4 control mice, **Fig. 1a**).

**Fig. 1:**
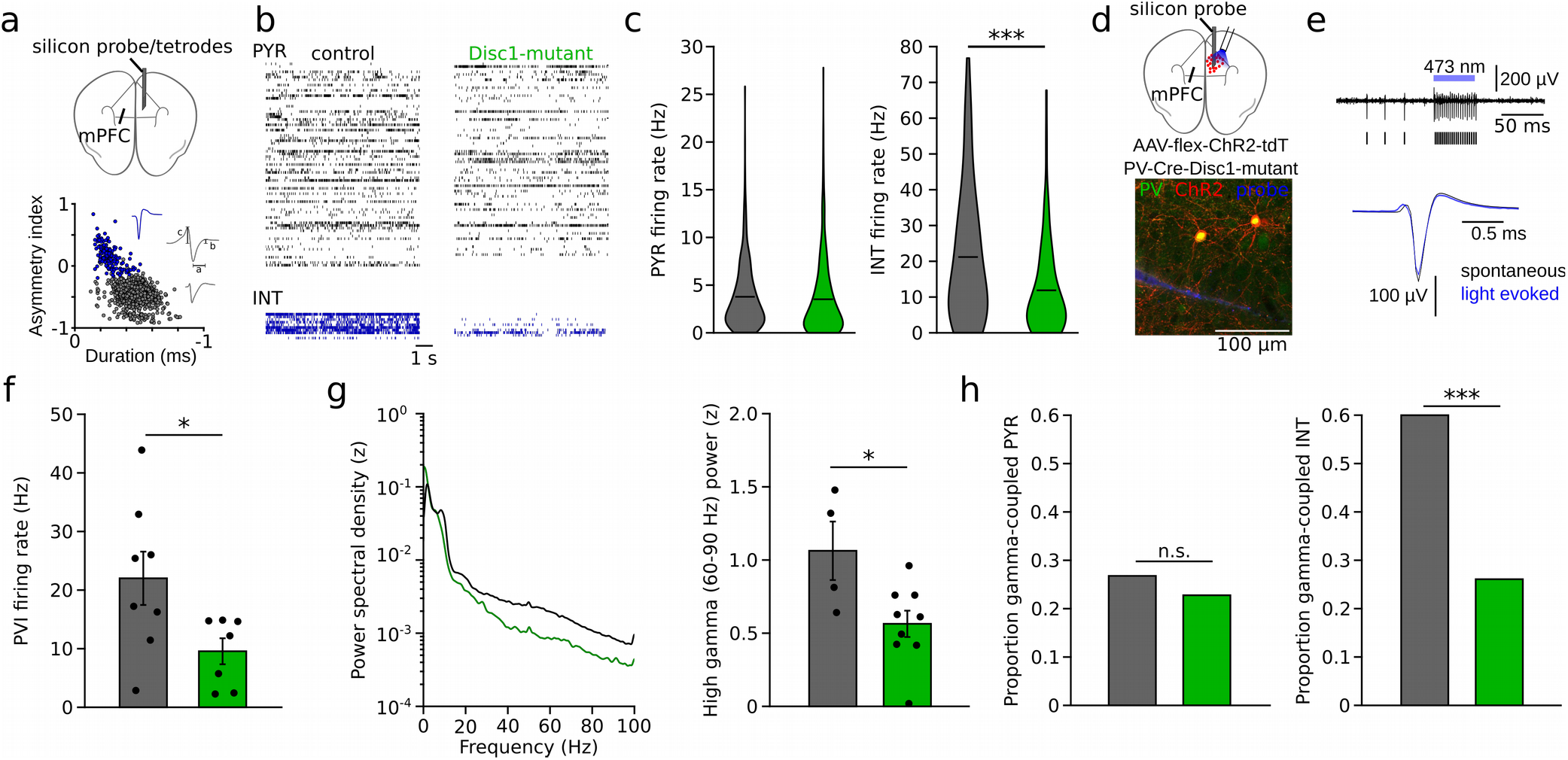
Reduced INT activity in the mPFC of Disc1-mutant mice. **a** Schematic of the recording configuration. Silicon probes or tetrode microdrives were targeted to the mPFC of freely moving mice. Bottom, identification of PYRs (grey) and INTs (blue) based on duration a and asymmetry index (b-c)/(b+c) of filtered action potential waveforms. Blue and grey insets show an average unfiltered spike of an INT and PYR, respectively. **b** Example raster plots of PYR and INT activity in a freely moving control (black) and Disc1-mutant mouse (green). Each line shows the time series of spikes of a single neuron. **c** Decreased spike rates of INTs but not PYRs in the Disc1-mutant mPFC (Welch’s tests). **d** Schematic of the experimental strategy to label PVIs with ChR2. **e** Optogenetic identification of PVIs during blue laser illumination in awake head-fixed mice. Top: 0.3-6 kHz-filtered recording of one channel of the silicon probe. Bars indicate spike times of the unit. Bottom, comparable waveforms of light-triggered and spontaneous spikes. **f** Reduced spontaneous spiking of otpogenetically identified PVIs in head-fixed PV-Cre-Disc1-mutant animals. Mann-Whitney U-test. Data points are individual PVIs. **g** Reduced gamma oscillation power in freely moving Disc1-mutant mice. Left, example of power spectral density determined in the mPFC of a control (black) and Disc1-mutant mouse (green). Right, summary plot of high-gamma (60-90 Hz) power (unpaired t-test). Data points represent individual mice. **h** Quantification of the proportion of PYR (left) and INT (right) significantly phase-coupled to high-gamma oscillations. Coupling was determined with Rayleigh’s test. Comparisons between Disc1-mutant and control were done with χ^2^-tests. *p<0.05, ***p<0.001.

We found a significant reduction in INT rates in Disc1-mutant mice (t(176)=3.686, p=0.0003, Welch’s test), while the firing rates of PYRs were unaltered (t(1263)=1.280, p=0.201, Welch’s test, **Fig. 1b,c**). Since neocortical INT types are highly diverse (Gupta et al., 2000, Druckmann et al., 2013), putative INTs identified by their spike shape might include a mixed set of GABAergic populations. To specifically assess the activity profile of identified INTs, we focussed on PVIs by crossing Disc1-mutant animals with mice expressing Cre-recombinase under the control of the PV-promotor (**Fig. 1d**, PV-Cre-Disc1-mutant). Stereotaxic infusion of adeno-associated viruses encoding channelrhodopsin-2 (ChR2) and the red fluorophore tdTomato (tdT) flanked by double-inverted loxP sites (AAV-Flex-ChR2-tdT) into the mPFC rendered PVIs sensitive to blue light and allowed optogenetic identification of PV-expressing cells by brief laser pulses during single-unit recording (~10 mW, 50 ms; **Fig. 1d,e**). ChR2-expression was highly specific for PVIs (**Supplementary Fig. 2**). Optogenetic identification experiments were performed in awake, head-fixed mice, running on a circular track (**Methods**). In the absence of light stimulation, the discharge rate of PVIs was significantly reduced in PV-Cre-Disc1-mutant mice (U(13)=9, p=0.016, Mann-Whitney U-test, **Fig. 1f**). The reduction in PVI activity in Disc1-mutant mice could not be explained by an altered speed modulation because Disc1-mice ran at comparable velocity as control mice (**Supplementary Fig. 2**). Furthermore, similar results of reduced PVI rates were obtained during recordings under ketamine anesthesia in a separate cohort of mice (**Supplementary Fig. 3**).

Consistent with the essential role of PVIs in the emergence and synchrony of gamma oscillations (Bartos et al., 2007, Cardin et al., 2009, Sohal et al., 2009), the power spectral density in the high-gamma frequency range (60-90 Hz) was significantly reduced to ~52% in freely moving Disc1-mutant mice (U(11)=4.0, p=0.019, Mann-Whitney U-test, **Fig. 1g**), while theta or low-gamma oscillations were not significantly affected (**Supplementary Fig. 4**). Moreover, the proportion of INTs phase coupled to high-gamma oscillations was reduced from 60% in control to ~25% in Disc1-mutant mice (χ^2^(160)=18.937, p<0.001, χ^2^-test), while the proportion of significantly gamma-coupled PYRs was unchanged (χ^2^(1033)=2.224, p<0.135, χ^2^-test, **Fig. 1h**). Thus, Disc1 mutation results in impaired discharge rates of INTs, including PVIs, and decoupling of INTs from fast network oscillations.

### Impaired spike transmission at PYR-INT synapses of Disc1-mutant mice

To assess whether reduced glutamatergic drive from local PYR might contribute to lower INT activity levels, we investigated synaptic interactions from PYRs onto INTs *in vivo* (**Fig. 2**). These connections can be captured by spike train cross-correlations in simultaneous recordings of large neuronal populations (English et al., 2017, **Fig. 2a-c**). PYR-INT connections were assessed by determining spike transmission probability from baseline-corrected cross-correlograms of PYR and INT spike trains (English et al., 2017, **Fig. 2b**). Monosynaptic excitatory connections were apparent as significant and sharp positive peaks in causal direction (i.e. spiking in the PYR preceding spiking in the INT at short latency, **Fig. 2b,c**). Among 7639 PYR-INT pairs tested, we detected 138 significant monosynaptic interactions. The probability of finding PYR-INT connections was significantly reduced in the Disc1 mPFC (p=0.001, chisquare test, data not shown). Moreover, cross-correlation analysis revealed significantly reduced spike transmission to ~59% of control levels in the mPFC of Disc1-mutant-mice (t(136)=2.937, p=0.004, Welch’s test; **Fig. 2c,d**). Furthermore, spike transmission at connections to optogenetically identified PVIs were reduced to ~42% (t(15)=2.653, p=0.029, Welch’s test, **Fig. 2e**).

**Fig. 2:**
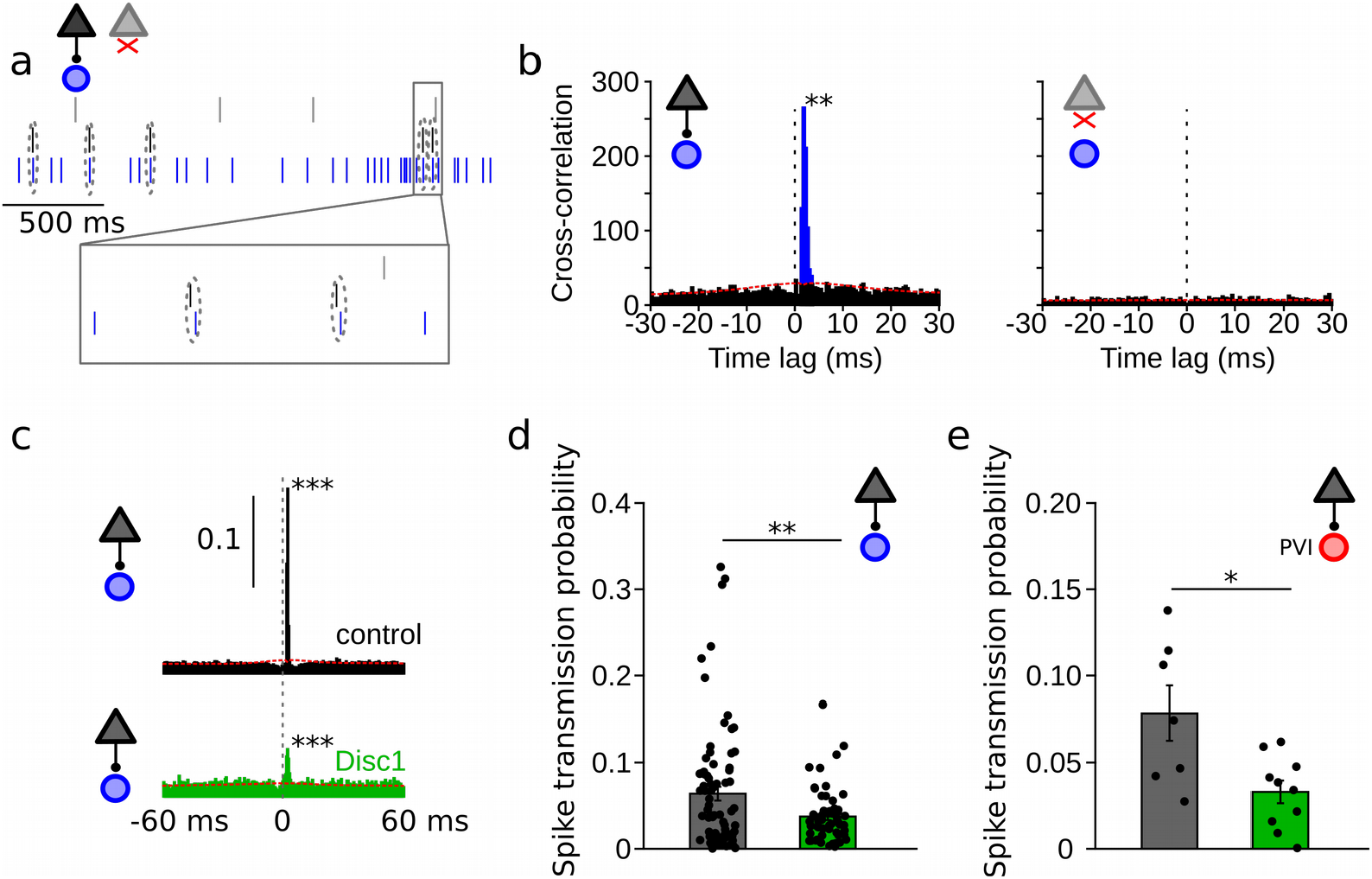
Reduced spike transmission at PYR-INT connections in the Disc1-mutant mPFC. **a** Short segment of spike trains of a PYR with (black) and without (grey) monosynaptic excitatory connection onto a GABAergic fast-spiking INT (blue). Note that spiking in the black PYR consistently occurs briefly before an action potential in the INT. **b** Cross-correlograms of the units shown in (a) reveal a sharp peak at 1-2 ms latency for the connected pair (left) but not for a non-connected pair of cells (right). Blue region shows short-latency spikes significantly exceeding the slowly co-modulated baseline (red dotted line, see **Methods**). **c** Example spike transmission probability histograms for a connected PYR-INT pair in control (black) and Disc1-mutant mPFC (green). A significant synaptic connection was detect in both cases (p<0.001). **d** Summary of spike transmission probabilities of all PYR-INT pairs in freely-moving mice (Welch’s test). **e** Summary of spike transmission probabilities of identified PYR-PVI connections in head-fixed mice (Welch’s test). Circles represent individual cells. *p<0.05, **p<0.01. Circles in d represent PYR-INT pairs and in e, PYR-PVI pairs.

We performed a series of analyses to decipher whether reduced spike transmission reflects a proper reduction in the excitatory drive to local interneurons, or whether it might (in part) be explained by altered network properties. First, we assessed the somatic distance between a presynaptic PYR and a postsynaptic INT in a subset of mice, in which recordings were performed with silicon probes (n=4 Disc1-mutant and 2 control mice, **Fig. 3a**). In these recordings, the position of the pre- and postsynaptic neurons could be estimated due to the known physical location of the electrodes with the largest spike waveform deflections. Consistent with previous reports, the strength of the spike transmission was inversely correlated with distance in both control (p=0.006) and Disc1-mutant mice (p=0.024, English et al., 2017, **Fig. 3b**). Connected pairs in Disc1-mutant and control mice were similarly spaced (t(83)=-0.206, p=0.837, Welch’s test, **Fig. 3c**). These results rule out changes in distance-dependent signaling properties or systematic differences in intersomatic distances of the recorded PYR-INT pairs as potential explanations of reduced spike transmission. We moreover consistently observed converging input of multiple (>1) PYRs onto one INT with no significant difference in the number of presynaptic partners between genotypes (U(64)=517.5, p=0.380, Mann-Whitney U-test, **Fig. 3d, left**). Similarly, PYRs of both groups made frequently contact with more than one postsynaptic INT, with no significant difference in the number of divergent targets (U(110)=1387, p=0.113, Mann-Whitney U-test, **Fig. 3d, right**), indicating intact overall PYR-INT network structure.

**Fig. 3:**
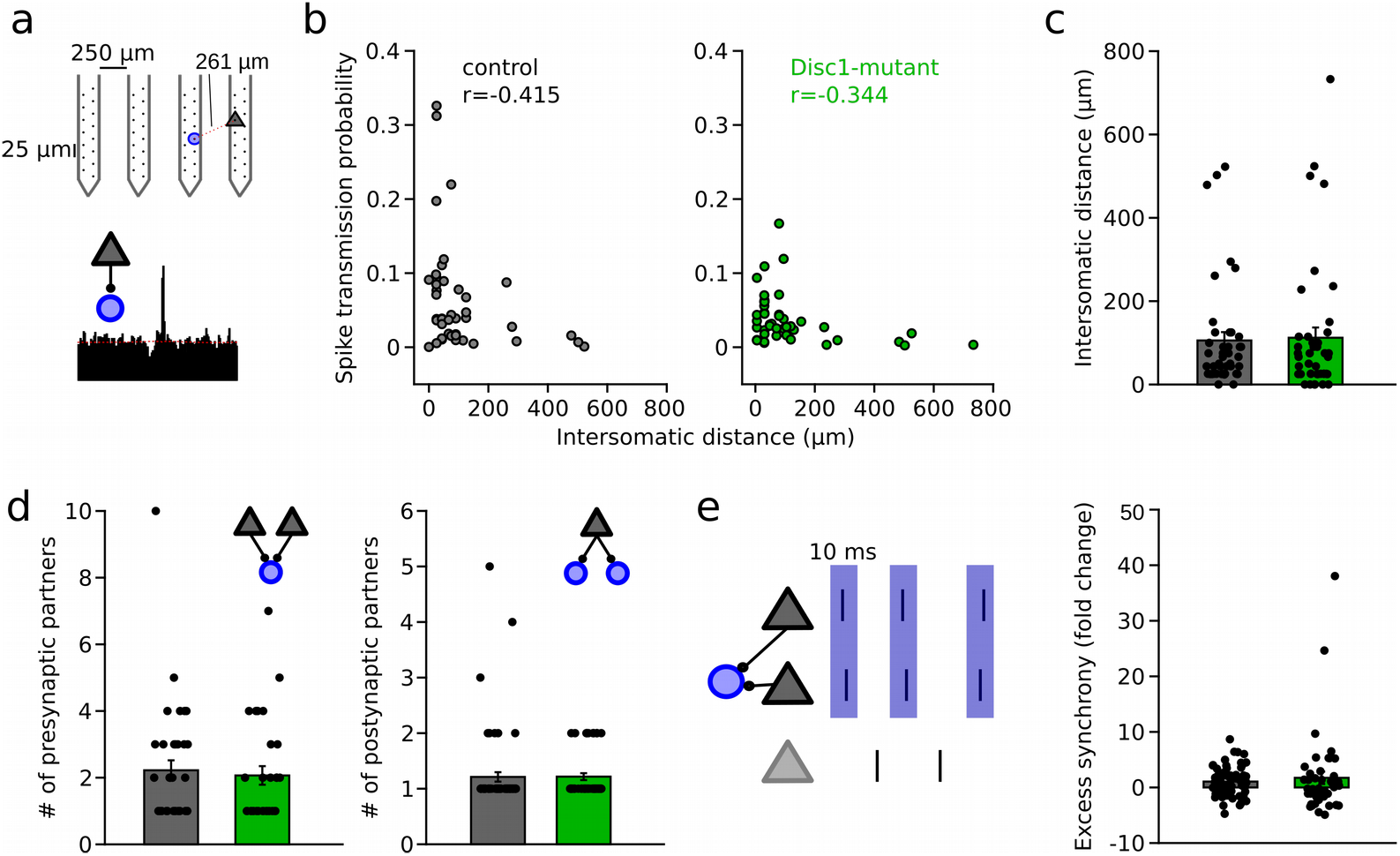
Unaltered network structure and presynaptic cooperativity of PYR-INT connections. **a** Example of distance calculation and corresponding cross-correlogram for a connected pair. **b** Summary of spike transmission probability as a function of intersomatic distance. Spike transmission inversely depends on intersomatic distance in both control (black) and Disc1-mutant pairs (green) (Spearman correlation coefficients). Circles are individual PYR-INT pairs. **c** Summary of intersomatic distances of all PYR-INT pairs (Welch’s test). **d** Quantification of the number of presynaptic (left) and postsynaptic partners (right) revealed comparable convergence and divergence, respectively (Mann-Whitney U-tests). **e** Left, schematic of synchronization of PYRs impinging on a shared postsynaptic INT. Right, summary of excess <10 ms synchrony between convergent PYRs (Welch’s test). Circles in b-e represent individual cells/cell pairs.

Second, synchronous presynaptic activity of PYRs with a common postsynaptic target has been demonstrated to boost spike transmission in the hippocampus (English et al., 2017). We extracted the pairwise synchronization of presynaptic PYRs with a common postsynaptic INT. Excess synchronization of convergent PYRs was not significantly different between Disc1-mutant and control mice (t(157)=-1.350, p=0.182, Welch’s test, **Fig. 3e**). These findings rule out defective synchronization of PYR as a mechanism of reduced spike transmission.

Third, hippocampal INTs show resonance membrane properties that boost spike transmission when postsynaptic spikes occur within a ~20-50 ms time interval (English et al., 2017). Consistent with reports from the hippocampus, neocortical PYR-INT connections displayed a ~3-fold boost of spike transmission probability in the resonance time window of 20-50 ms, similar to the gain observed in CA1 (English et al., 2017, **Supplementary Fig. 5**). Moreover, identified PVIs showed ~2-fold resonance boost of spike transmission (t(10)=-3.406, p=0.019, paired *t-*test, **Supplementary Fig. 5**). However, the magnitude of the resonance gain reached similar levels for PYR-INT pairs in Disc1-mutant mice (t(124)=1.089, p=0.278, Welch’s test, **Supplementary Fig. 5**). In identified PVIs, resonance boosting of spike transmission was even significantly higher in Disc1 compared to controls (U(14)=11, p=0.022, Mann-Whitney U-test, **Supplementary Fig. 5**). Finally, reduced spike transmission in Disc1-mutant mice was also evident when we only considered spikes in the resonance time window (t(124)=2.660, p=0.009, Welch’s test, **Supplementary Fig. 5**). Jointly, these data suggest that while INTs in the Disc1-mutant mPFC display intact convergence, responses to synchronous input and resonance properties, they respond less to the action of local glutamatergic synaptic transmission.

### Impaired neuronal assembly structure in the Disc1 mPFC

Previous experimental and simulation studies suggested that an important function of GABAergic inhibition might be to decorrelate synchronous activity of pyramidal cells imposed by shared inputs (Renart et al., 2010, Tetzlaff et al., 2012, Sippy and Yuste, 2013). Theoretical work further emphasized that feedback inhibition within a circuit is particularly effective at constraining pairwise spike train correlations (Tetzlaff et al., 2012). We therefore hypothesized that reduced inhibitory activity levels might affect PYR coactivity in the Disc1-mutant mPFC. In the Disc1 mPFC, PYRs indeed displayed stronger pairwise synchronization at 10 ms time scale (t(33012)=-5.312, p=1.1*10^-7^, n=14524 control and 18490 Disc1 PYR pairs, this analysis considered all PYRs, **Fig. 4a**).

**Fig. 4:**
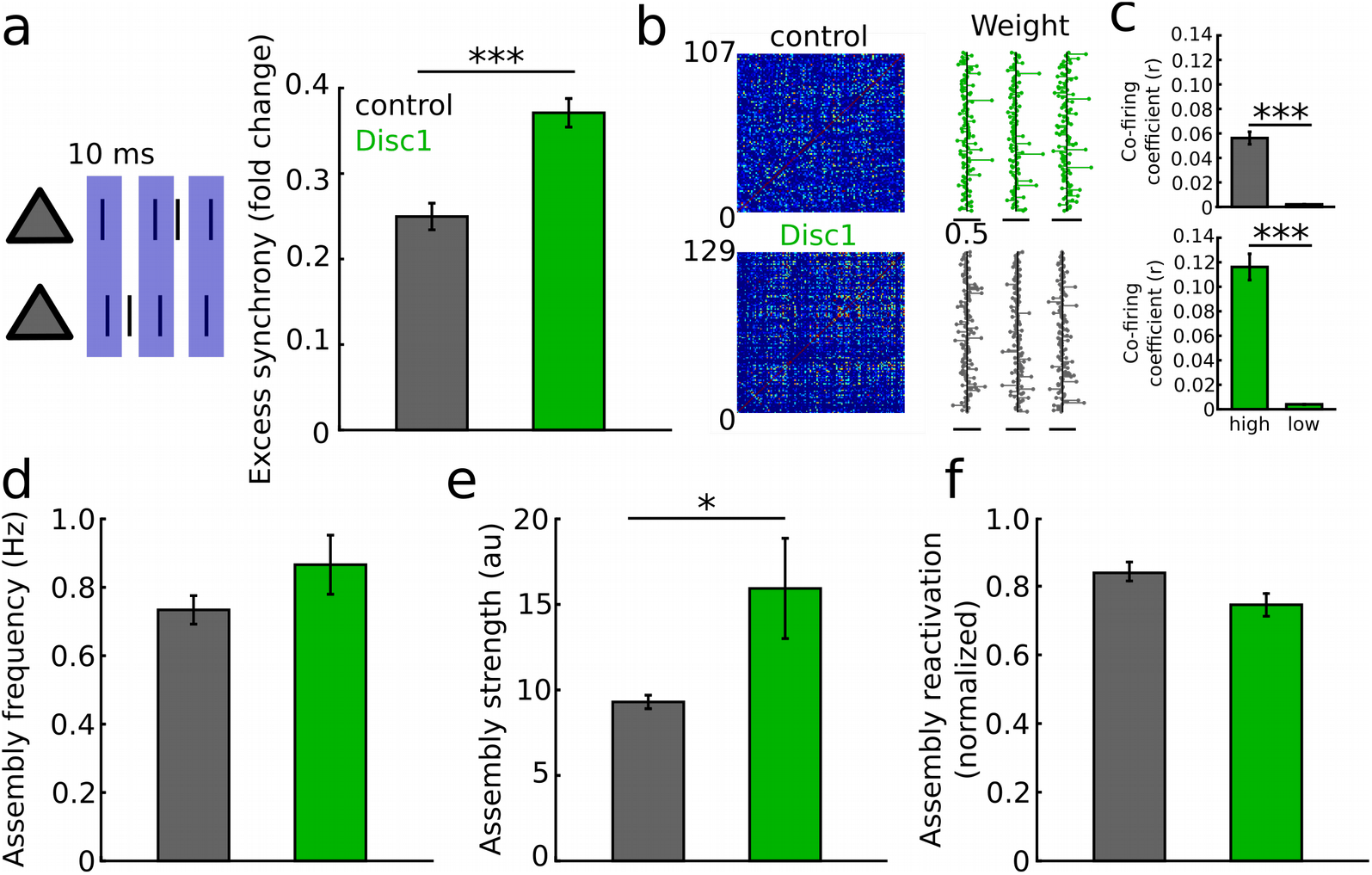
Enhanced synchronization and assembly activations in the Disc1-mutant mPFC. **a** Left, schematic of synchronization of PYRs. Right, summary of excess <10 ms synchrony (Welch’s test). **b** Detection of assembly patterns. Co-active patterns were extracted from spike covariance matrices (left) to retrieve assembly weight vectors (right). Data show examples from one control and Disc1-mutant mouse. **c** Neurons with high weight in an assembly pattern show stronger cofiring than neurons with low contribution (Welch’s tests). **d** Assemblies activate on average at the same frequency in both genotypes. **e** Assembly activation strength is enhanced in Disc1-mutant mice. **f** Control and Disc1-mutant assemblies show comparable stability (comparison for the mean expression strength during the first and second half of the recording). Data in **d-e** are from 101 and 110 assembly patterns in control and Disc1-mutant mice, respectively (Welch’s tests). *p<0.05, ***p<0.001.

Groups of co-active cells forming neuronal assemblies are thought to be a hallmark of cortical processing (Buzsáki, 2010, El-Gaby et al., 2021). To directly assess the expression of neuronal assembly patterns beyond pairwise comparisons, we retrieved co-active firing motifs using a step-wise extraction procedure relying on principal and independent component analysis of binned spike trains of simultaneously recorded PYRs (van de Ven et al., 2016, El-Gaby et al., 2021, **Fig. 4b**). Using this procedure, we detected in total 101 assembly patterns in control mice (n=15 sessions) and 110 patterns in Disc1-mutant mice (n=18 sessions), with one assembly pattern found per 4.9 ± 0.4 and 5.2 ± 0.5 PYRs in control and Disc1 mPFC, respectively (t(31)=-0.463, p=0.647). Neurons with large weight in a detected assembly pattern showed more pronounced co-firing than neurons that contributed less to a given pattern (**Fig. 4c**), corroborating the validity of the assembly detection method. Spontaneous assembly patterns activated at comparable rates in control and Disc1-mutant mice (t(209)=-1.371, p=0.172, **Fig. 4d**). However, consistent with enhanced pairwise co-firing, the assembly activation strength was significantly larger in Disc1-mutant mice (t(209)=-2.232, p=0.028, **Fig. 4e**). Furthermore, we observed a trend towards reduced stability of assembly patterns over the course of the recording (t(179)=1.962, p=0.051, **Fig. 4f**). These data jointly indicate altered neuronal coactivation and assembly structure in Disc1-mutant mice.

## Discussion

Despite the broad recognition that fast-spiking INTs, particularly PVIs, contribute to neuronal network synchronization at gamma frequencies (Bartos et al., 2007, Cardin et al., 2009, Sohal et al., 2009) show task-dependent tuning during working memory (Kim et al., 2016a, Lagler et al., 2016), and are required for the proper execution of working memory (Murray et al., 2011, Murray et al., 2015), information is still lacking on PVIs’ activity in the prefrontal cortex of working memory-deficient Disc1-mutant mice. A major consequence related to this gap of knowledge is our limited understanding of the underlying pathophysiology, which prevents the development of new therapies for mental diseases. Here, we provide a first *in vivo* electrophysiological characterization of fast-spiking INT and PVI activity in Disc1-mutant mice. Extending upon previous electrophysiological Disc1 studies (Sauer et al., 2015, Chini et al., 2020, Kaefer et al., 2019, Delevich et al., 2020), we show that the average discharge rates of INTs are reduced *in vivo* by ~44% whereas PYR activity is unaltered (**Fig. 1b,c**).

Related to the reduced mean PVI activity, we detect a ~41% reduction in the magnitude of spike transmission probability at PYR-PVI connections and a reduced probability of finding connected PYR-INT pairs in Disc1-mutant mice (**Fig. 3d,e**). Spike transmission is an indirect estimation of synaptic connections onto INTs (English et al., 2017). Control analyses demonstrated that the weaker spike transmission is unlikely to be caused by network effects such as altered convergence or intersomatic distance between the connected cells, suggesting that it reflects a genuine reduction in excitatory drive at PYR-INT connections. Given the lower spike transmission probability in Disc1-mutant mice, it is, however, likely that the number of detected pairs might have been under-sampled in Disc1-mutant mice. We propose that reduced drive to INTs induces a chain of interrelated functional consequences in the inhibitory network. Diminished recruitment of PVIs by local glutamatergic inputs will result in lower PVI firing rates and weaker PVI-mediated synaptic inhibition. Lower synchrony of PVIs might consequently result in the observed reduction in gamma power in Disc1-mutant mice (Kim et al., 2016b; Wang and Carlén, 2012). A gamma oscillation dysfunction in the mPFC might contribute to working memory deficits in Disc1-mutants. This proposal fits to reports of high-gamma activities occurring during the execution of working memory in mice and monkeys (Lundqvist et al., 2016; Yamamoto et al., 2014).

One might expect that reduced PVI activity leads to enhanced PYR discharge rates due to reduced perisomatic inhibition (Courtin et al., 2014). However, we did not observe significant changes in the average activity or gamma-phase relationship of PYRs. Since perisomatic inhibition generates large GABA_A_ receptor-mediated conductances at unitary GABAergic connections (Jouhanneau et al., 2018), the compound inhibitory conductance emerging from the convergence of active GABAergic perisomatic inputs onto a single PYR might be sufficiently large to control the activity of PYR populations. This hypothesis fits to previous studies showing that pharmacogenetic silencing of hippocampal PVIs or the elimination of PVI synaptic output with tetanus toxin had no effect on the expression of the activity marker cfos (Murray et al., 2011; Stefanelli et al., 2016).

In addition to effects on firing rate, it has been argued that inhibition might counteract excessive synchronization of PYRs be effectively decorrelating shared synaptic inputs (Renart et al., 2010, Tetzlaff et al., 2012, Sippy and Yuste, 2013). In line with this hypothesis, chemogenetic silencing of PVIs enhanced pairwise correlations among cortical PYRs (Hamm et al., 2017). Supporting the synchronization-limiting role of feedback inhibition, we found increased spike correlations among cortical PYRs in Disc1 mutant mice. This observation generalized to identified neuronal assembly patterns, which showed stronger expression strengths. The expression strength of neuronal assemblies as evaluated with the method chosen here reflects the number of coative cells, their synchronization and firing rate. Importantly, we chose a time window of 25 ms to detect assembly patterns, which is within the integration time of cortical PYRs (Koch et al., 1996). Active assemblies within this integration window are thus expected to efficiently affect downstream reader neurons (Buzsáki, 2010).

Disturbed cell assembly expressions have been identified in the chronic ketamin and Df(16) models of schizophrenia (Hamm et al., 2017). Notably, in those models, stability of assembly patterns rather than their expression strength were affected. These data imply that while the nature of the assembly dysfunction might differ between distinct models, altered assembly expressions might be a common final path underlying cognitive distrubances in schizophrenia.

## Methods

### Animals

Disc1-mutant, PV-Cre-Disc1-mutant, PV-Cre and wild type control mice (Bl6/J) were maintained on a 12h dark-light cycle with free access to food and water. Disc1-mutant mice were a gift from J. Gogos (Koike et al., 2006). At the start of the experiment, animals were at least 6 weeks old. Mice of both sexes were used in this study. All experiments were performed in agreement with national legislation.

### Working memory test

Mice were food-restricted to reach 85-90 % of their free-feeding body weight. After habituation to the T-maze for two days and two days of shaping, mice were trained to alternate on consecutive runs. In the sample run, one of the two target arms was blocked and the mice received a reward at the end of the open target arm. After return to the start box, a sample run was conducted during which both target arms were accessible. Mice were trained with 10 daily runs until a criterion of >70% correct was reached. After three consecutive days above criterion, testing commenced with a 15 sec delay between the sample and the match run.

### Microdrive implantation and electrophysiological recording

Mice were anesthetized with isoflurane in oxygen (induction: 3%, maintenance: 1-2%) and placed on a heating pad. Analgesia was achieved by subcutaneous injection of buprenorphine (0.05-0.1 mg/kg body weight). A four-shank silicon probe (model P-1, Cambridge Neurotech) mounted on a microdrive (NanoDrive, Cambridge Neurotech) was implanted in rostro-caudal orientation with the most anterior shank positioned at ~2 mm anterior from bregma and 0.35-0.4 mm lateral to the midline. A pair of stainless-steel screws (M1) was inserted into the bone over the cerebellum and connected to the ground and reference leads of the electrode interface board. Additional screws were inserted into the parietal and contralateral frontal bone for further stability. In a subset of mice, microdrives containing eight tetrodes build from 12.5 μm tungsten wire were targeted to the mPFC, using the following coordinates: 1.8-2 mm anterior of bregma, 0.45 mm lateral of the midline, and 1.8-2.8 mm below bregma. The microdrive was fixed to the skull with dental cement (SuperBond). After surgery, analgesia was provided for two days with buprenorphine (subcutaneous injections every 6h during daytime and in the drinking water over night) and carprofen (4-5 mg/kg body weight, subcutaneous injection every 24 h for two days). About one week after surgery, wide-band neural signals (0.1 Hz-7.5 kHz, 32-64 channels) were recorded with tethered RHD2000 amplifiers (Intan Technologies) at 30 kHz sampling frequency while animals were awake in their home cage. Data acquisition was performed with the OpenEphys GUI. After the recording, the microdrive was lowered by 100-200 μm to obtain access to a new set of units on the next recording day. Up to five recording sessions were performed with each animal. After completion of the recordings, the mice were deeply anesthetized by an interaperitoneal injection of urethane (2g/kg body weight) and transcardially perfused with ice-cold phosphate-buffered saline (~10 ml) followed by freshly prepared ice-cold 4% paraformaldehyde (50-100 ml). In a subset of animals, electrolytic lesions of the recording sites were done with DC stimulation (10-20 V) applied to a subset of electrodes. After post-fixation overnight, 100 μm sections were cut with a vibratome (Leica VT1200 S) and stained with DAPI. Epifluorescence microscopy was used to assess the location of the recording sites. Recordings from the prelimbic, cingulate and infralimic cortex were included and pooled for analysis.

### Viral injections

Animals were anesthetized with isoflurane as described above. AAV1-flex-ChR2-Tdtomato (Addgene, titer 2*10^12^/ml, 0.5 μl) or AAV1-flex-ChR2-mCherry (Charite Vector core, titer 4*10^u^/ml, 0.5 μl) was slowly infused into the right or left mPFC at two anterio-posterior locations (1.5 and 2 mm anterior from bregma) using thin glass pipettes fabricated with a microfilament puller (Flaming Brown). After viral injection, the pipette was kept in place for 5 min to ensure efficient virus diffusion into the tissue. Acute recordings commenced >2 weeks after virus injection.

### Acute recordings

For recordings under anesthesia, the mice were anesthetized with isoflurane and a craniotomy (~2 mm wide) was performed above the right mPFC. After removal of the dura, the craniotomy was sealed off with dura gel (Cambridge Neurotech) and a custom 3D-printed head bar with a circular ring around the craniotomy site was fixed to the skull with dental cement. The animal was then immediately transferred to the recording station while anesthesia was maintained throughout the recording session with intraperitoneal injections of ketamine and xylazine (initial dose: 100 and 13 mg/kg body weight, respectively, topped up by 10-20% every 20-40 min). For awake recordings, a steel head plate was implanted on the skull and the animals were allowed to recover from head plate implantation for three days. For habituation to head fixation, the mice were briefly sedated with isoflurane and head-fixed such that they could comfortably stand on a circular Styrofoam weal. A virtual reality (circular track, length 1.5 m, visual cues placed outside the arena) was constructed with open-source 3D rendering software (Blender) and was projected on five computer screens surrounding the head-fixation setup (Schmidt-Hieber and Häusser, 2013). Over subsequent days, mice were accustomed to head fixation by daily increasing the time of head-fixation until the animals appeared calm and traversed the circular maze reliably. Once the animals were habituated, a craniotomy was performed as described above. Carprofen was injected subcutaneously on the day of the surgery. A 4-shank silicon probe (Cambridge Neurotech) coated with fluorescent marker (DiD) was slowly (~5-10 μm/s) lowered to the mPFC (935-1758 μm below brain surface). Wide-band neural signals were recorded at 30 kHz sampling with a 64-channel amplifier (Intan Technologies) connected to an USB acquisition board (OpenEphys). Laser light (473 nm, ~10 mW intensity at the fiber tip, 50 ms pulses at 0.1 Hz) was delivered through a 200 μm optical fiber glued to the silicon probe. Afterwards, the silicon probe was slowly retracted and the animals were transcardially perfused with ~20 ml phosphate-buffered saline followed by ~30 ml of 4% paraformaldehyde. After post-fixation overnight, 100 μm-thick frontal sections of the mPFC were cut and stained with rabbit-anti-PV antibody (1:1000, Swant) and DAPI. The location of the silicon probe and the immunostaining were visualized with a laser-scanning microscope (LSM 710, Zeiss). Recordings in the prelimbic and cingulate cortex were included in this study and pooled for analysis.

### Single-unit isolation

Single unit clusters were isolated from bandpass-filtered raw data (0.3-6 kHz) using MountainSort (Chung et al., 2017). Putative single-unit clusters that fulfilled quality criteria of high isolation index (>0.90) and low noise overlap (<0.1) were kept for manual curation, during which only clusters with a clear refractory period and clean waveform shape were saved for further analysis. In case of two clusters with similar waveforms, cross-correlation was used to assess whether clusters had to be merged. Individual tetrodes or shanks of a silicon probe were clustered separately. Isolated units were separated in excitatory and inhibitory neurons based on trough-to-peak duration and asymmetry index (Sirota et al., 2008) using *k*-means clustering. To analyze physical distances between units, the channel on the silicon probe with the largest negative amplitude deflection was defined as the location of the unit and the absolute inter-somatic distance was calculated.

### Optogenetic tagging

To detect ChR2-expressing PVIs, we applied 50 ms pulses of blue laser light. The firing rate during the last 25 ms of the pulse was calculated and averaged for all pulses. Next, we created a random spike train of the unit by shuffling the interspike-intervals and calculated the average spike frequency during the second half of the light pulse. The shuffling was repeated independently 1000 times. A unit was considered significantly light-sensitive if the actual rate during light delivery exceeded the 99^th^ percentile of that of the shuffled distribution.

### Spectral analysis and spike-field coupling

For power spectral density analysis, the raw traces were z-scored and power was measured in a sliding window (length: 1s) with fast-Fourier transforms (zero-padding: 10 s). Power was averaged across five randomly chosen channels for each mouse.

To assess phase-coupling to the local field potential, the raw local field potential trace was filtered in the high-gamma range (60-90 Hz). The instantaneous phase at each time of spiking of a given neurons was extracted using the Hilbert transform. Significant coupling was tested with Rayleigh’s test for circular uniformity applied to the resulting phase angles. The coupling depth of significantly coupled cells was determined using the pairwise phase consistency defined as the average pairwise circular distance (Tamura et al., 2016).

### Spike transmission analysis

To detect monosynaptic excitatory interactions between PYRs and INTs we utilized cross-correlation methods (English et al,. 2017). Cross-correlations between spike trains (0.4 ms bins, maximal time lag: 50 ms) were calculated if both units fired at least 500 spikes during the recording interval. Criteria for a significant monosynaptic interaction were a peak in the monosynaptic time window (0.8-2.8 ms following the spike in the PYR) significantly exceeding the co-modulated baseline and the peak in anti-causal direction (i.e. INT-PYR, −2 to 0 ms). The baseline *b* was obtained by convolving the raw crosscorrelogram with a partially hollowed Gaussian function (hollow fraction: 0.6, standard deviation: 10 ms). The Poisson distribution with continuity correction was used to estimate the probability of the observed magnitude of cross-correlation in the monosynaptic bins (*P_syn_*),

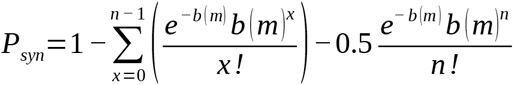

Similarly, we estimated the probability of the observed count in the monosynaptic bins of the cross-correlogram being larger than the count in anticausal direction (canticausal) using the Poisson distribution with continuity correction,

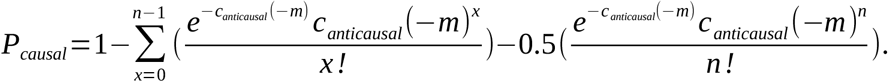

Following optogenetic ground truth data obtained in the hippocampus, a pair was marked as connected if P_syn_<0.001 and P_causal_<0.0026 (English et al., 2017). Spike transmission probability was defined as the spiking in the monosynaptic window exceeding *b* normalized by the number of presynaptic spikes.

### Presynaptic synchronization and resonance analysis

To assess presynaptic synchronization of PYRs with shared postsynaptic partner, we quantified the proportion of spikes of the two neurons that co-occurred within a time window of 10 ms. The data were then compared to a random distribution generated by randomly shifting the spike of both units between 1 ms and 30 s (1000 iterations). Excess synchrony was quantified by subtracting the average synchrony of the random distribution from the measured synchrony and dividing the result by the standard deviation of the random distribution (English et al., 2017).

To quantify resonance properties, presynaptic spikes that occurred at times when the postsynaptic neuron had fired 20-50 ms before the reference spike (resonance time window) and neither the pre-nor the postsynaptic unit had fired between the end of the resonance time window and the reference spike were isolated. Spike transmission probability was computed separately for these extracted resonance spikes and compared to the transmission probability obtained for all presynaptic spikes.

### Detection of cell assemblies

The spike trains of PYRs were binned in 25 ms bins and normalized by z-scoring. The number of assembly patterns was detected from the the binned spike train matrix using the Marčenko-Pastur law (van de Ven et al., 2016, El-Gaby et al., 2021). The Marčenko-Pastur law states that a covariationmatrix constructed from statistically independent random variables (such as neurons that do not co-fire) gives eigenvalues below a critical value (Lopes-dos-Santos et al., 2013). If neuronal firing occurs correlated with each other (as would be the case for assemblies), eigenvalues above the critical limit should exist. The number of eigenvalues thus indicates the number of assembly patterns. We determined the eigenvalue limit *l* as

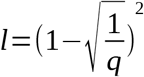

with

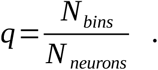

Using the fastICA algorithm of scikit.learn, we extract the number of independent components given by the eigenvalues above *l.* The resulting components are the weight vectors of each assembly. Since the orientation of independent components is arbitrary, each vector was oriented to have the largest deflection in positive direction and was further scaled to unit length (van de Ven et al., 2016). Assembly neurons were defined as those cells with a weight exceeding 2x the standard deviation of the pattern vector. To assess cofiring of neurons with large weight, Pearson’s *r* was measured for the binned spike trains of those neurons and compared to the correlation values among all other PYRs.

### Reconstruction of assembly activations over time

To obtain the activation strength *A* of assembly patterns, the weight vectors were projected on smoothed spike trains of all simultaneously recorded pyramidal neurons as

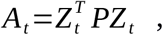

where *Z* is the smoothed and z-scored spike train of each neuron obtained by convolving the spike trains with a Gaussian function, *P* is the outer product of the respective weight pattern, and *T* is the transpose operator. Significant pattern activations where defined as threshold crossings of 5 (van de Ven et al., 2016). Average activation strength of a given pattern was calculated for all data points above that threshold. To measure assembly reactivation, the pattern detection was applied to the first half of the recording. Then, the expression strength over time was separately computed for the first and second half of the recording. Reactivation strength was measured as the average above-threshold strength of each pattern during the second half divided by the strength during the first half.

### Statistical analysis

Comparisons of two groups were performed with a two-sided Welch’s test if the number of observations was >30 in each group. For n<30 in each group, data were compared by two-sided student or Welch’s t-tests in case of comparable or different standard deviations, respectively, in case that data passed the normality test (Shapiro-Wilk test), otherwise a two-sided Mann-Whitney U-test was used. Paired comparisons were performed with paired t-test (two-tailed). Connection probabilities were compared with a χ^2^-test. All analysis (except for initial spike sorting, see above) including statistics were performed using Python2.7.

## Acknowledgements

We thank Karin Winterhalter and Kerstin Semmler for technical assistance and Sebastian Kugler for contributing to pilot experiments. This work was supported by the Deutsche Forschungsgemeinschaft (FOR2143-2, BA1582/2-2 M.B.; SA3609/1-1, JFS), the European Research Council Advanced Grant (ERC-AdG 787450, M.B.), and the Excellence Initiative of the German Research Foundation (Brain-Links Brain-Tools, M.B.).

## Author contributions

J-F.S and M.B. conceived the study, designed the experiments and wrote the manuscript. J-F.S. performed experiments and analyzed the data.

