## Supplementary Information for "*Disrupted-in-Schizophrenia-1* is required for proper pyramidal cell-interneuron communication and network dynamics in the prefrontal cortex"

Supplementary Materials:

Supplementary Figure 1-5

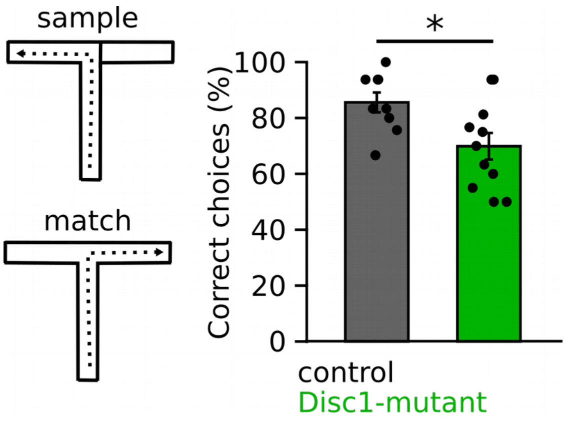


**Supplementary Figure 1: Working memory deficits of Disc1-mutant mice.** Working memory was assessed in a delayed-non-match-to-place paradigm, in which mice needed to remember the location of the previously visited arm during the match run. The summary bar graphs show the performance of Disc1-mutant (green) and control mice (black). T(18)=2.541, p=0.020, unpaired t-test. Symbols represent individual mice.

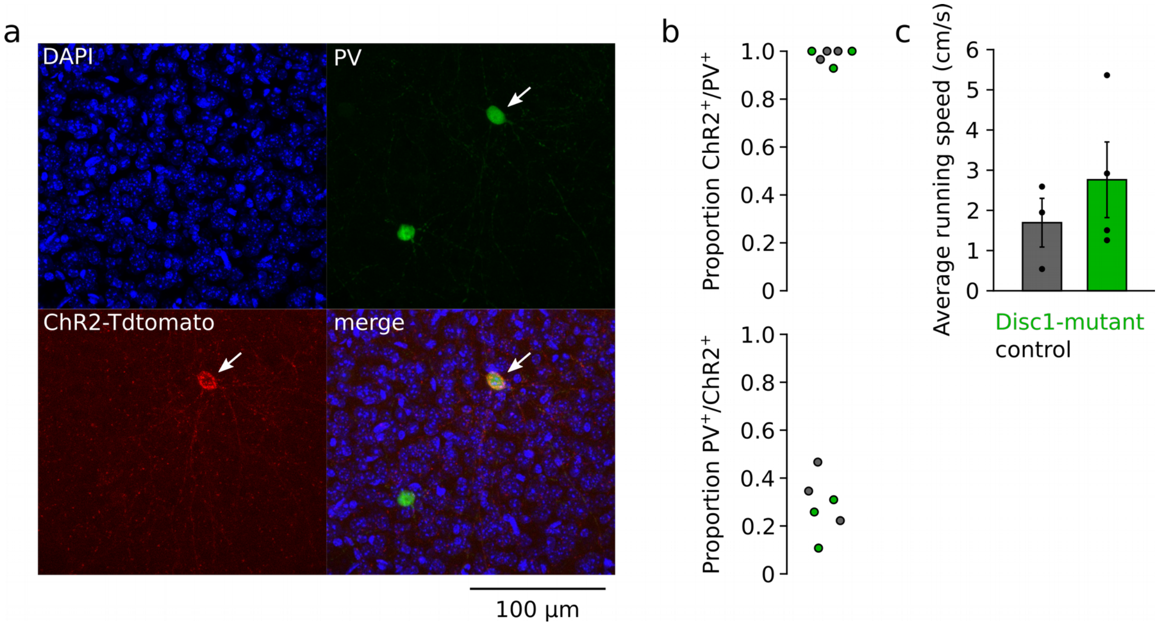


**Supplementary Figure 2: Specific expression of ChR2 in PVIs and comparable running speeds during head-fixation**. **a** Immunostaining against PV in a ChR2-infected mouse was used to determine colocalization of PVI and ChR2. **b** Quantification of colocalization from confocal images revealed high specificity of the viral approach (top) with the majority of ChR2-positive cells coexpressing PVI. On average ~30% of PVI were labelled by ChR2 with our approach (bottom). Circles are averages of individual Disc1-mutant (green) and control mice (black)**. c** Quantification of average running speed during head-fixation of mice exposed to a circular maze (**Methods**) revealed no difference between Disc1-mutant (green) and control mice (black). U(5)=4.0, p=0.298, Mann-Whitney U-test. Symbols represent individual mice.

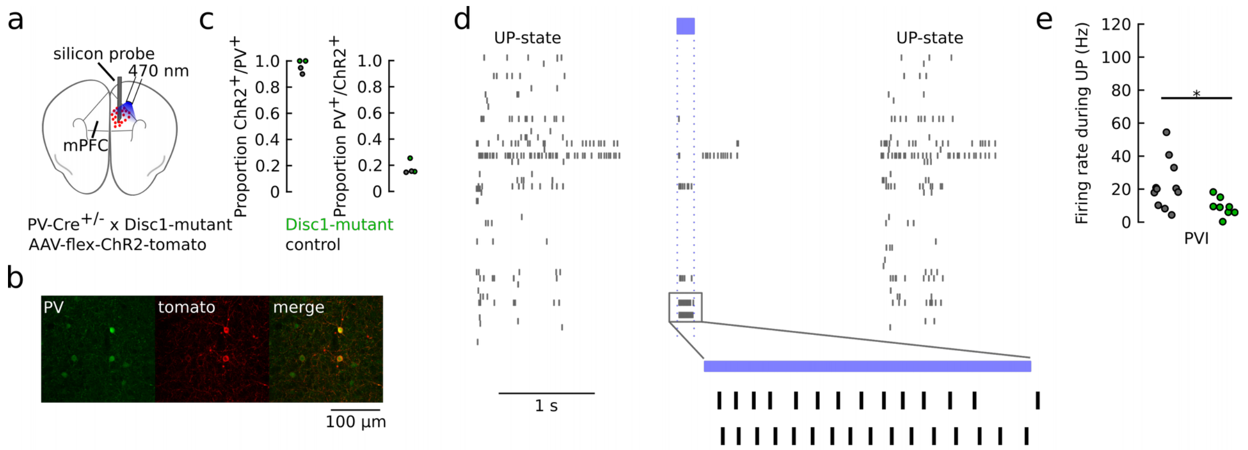


**Supplementary Figure 3: Reduced firing rate of optogentically identified PVIs during UP-states under ketamine anesthesia. a** Schematic of the recording strategy. The mPFC of PV-Cre-Disc1-mutant-mice was stereotaxically injected with AAV-flex-ChR2-tdT. A silicon probe was acutely inserted into the mPFC of ketamine-anesthetized mice. **b** example co-labelling of PV antibody staining (green) and tdT-expression in the mPFC. **c** quantification of tdT-ChR2-coexpression revealed that ChR2-expression was highly specific for PVIs. Each dot represents one control (black) or Disc1-animal (green). N=64 ChR2-positive and n=349 PV-positive cells were used for coexpression analysis. **d** Spike trains of simultaneously recorded neurons from one mouse under ketamine anesthesia. Firing is characterized by dense activity during neocortical UP-states intermingled with periods of relative silence. Blue laser light was applied through an optical fiber to excite ChR2-positive PVIs for optogenetic identification (blue bar). **e** Average UP-state firing rate of optogenetically identified PVIs (control: n=11, Disc1: n=8, 2 mice each, p(17)=2.419, p=0.027, unpaired t-test. *p<0.05.

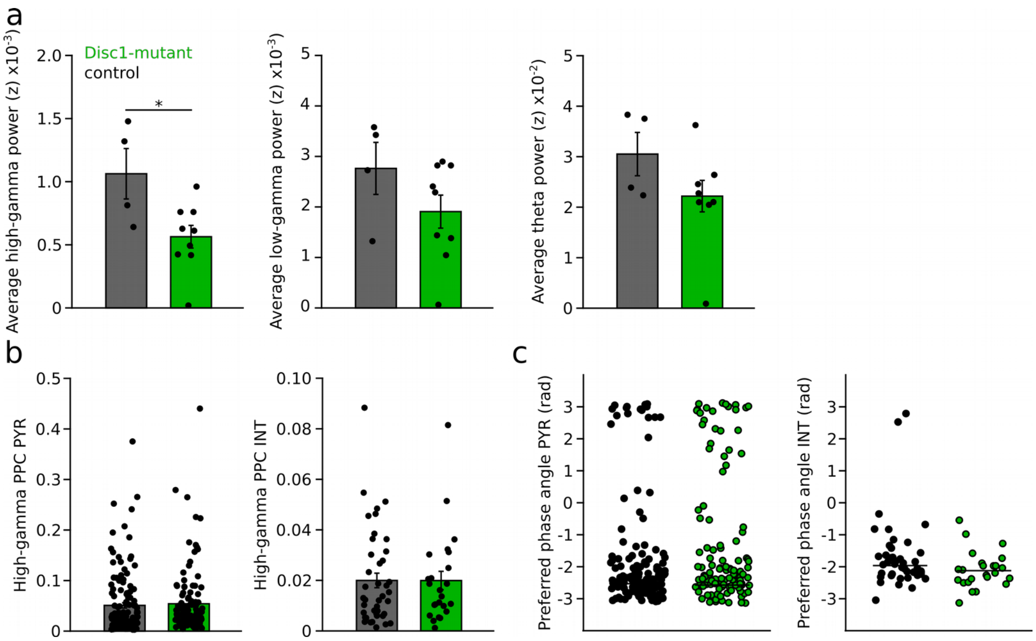


**Supplementary Figure 4: Phase-coupling of PYRs and INTs to local field potential oscillations. a** Power spectral density analysis revealed a reduction in the high-gamma band (60-90 Hz, right, U(11)=4, p=0.019) or and a trend towards a reduction in the low-gamma band (20-40 Hz, middle, U(11)=8, p=0.071) but not in the theta frequency range (left, 6-12 Hz, U(11)=10, p=0.124, Mann-Whitney U-tests). **b** Phase-coupling analysis of PYRs and INTs to high-gamma oscillations revealed comparable coupling depth, quantified as pairwise phase consistency (PPC; PYR: t(253)=-0.414, p=0.679, Welch's test, INT: U(64)=480, p=0.377, Mann-Whitney U-test) **c** PYRs and INTs discharged at similar preferred phase angles during high-gamma oscillations (PYR: p=0.612, INT: p=0.314, angle permutation test).

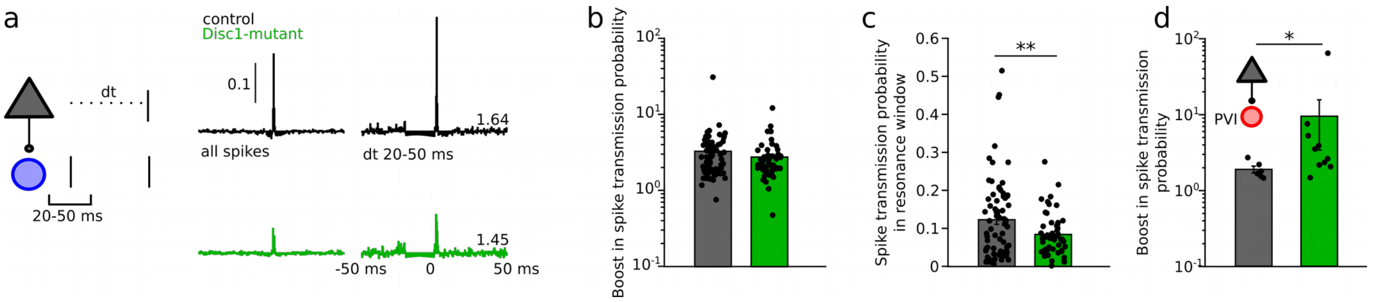


**Supplementary Figure 5: Increased resonance boosting at PYR-fast-spiking INT connections in Disc1-mutant mice. a** Left, schematic illustrating the quantification of spike transmission probability *in vivo* during a defined inter-spike interval. Right, examples of transmission gain for spikes at a resonance time window of 20-50 ms. Values indicate fold gain in the resonance time window. **b** Summary of the boost in spike transmission probability for PYR-INT pairs (Welch's test). **c** Summary of spike transmission probability in the resonance time window (Welch's test). **d** Summary of resonance boosting at PYR-PVI connections recorded in awake head-fixed mice (U(15)=11, p=0.022, Mann-Whitney U-test). Symbols are individual connections.
